# Tenascin-C potentiates Wnt signaling in thyroid cancer

**DOI:** 10.1101/2024.11.04.621959

**Authors:** Heather A. Hartmann, Matthew A. Loberg, George J. Xu, Anna C. Schwarzkopf, Sheau-Chiann Chen, Courtney J. Phifer, Kailey Caroland, Hua-Chang Chen, Diana Diaz, Megan L. Tigue, Amanda B. Hesterberg, Jean-Nicolas Gallant, Sophia M. Shaddy, Quanhu Sheng, James L. Netterville, Sarah L. Rohde, Carmen C. Solórzano, Lindsay A. Bischoff, Naira Baregamian, Paula J. Hurley, Barbara A. Murphy, Jennifer H. Choe, Eric C. Huang, Fei Ye, Ethan Lee, Vivian L. Weiss

## Abstract

Tenascin-C (TNC) is a secreted extracellular matrix protein that is highly expressed during embryonic development and re-expressed during wound healing, inflammation, and neoplasia. Studies in developmental models suggest that TNC may regulate the Wnt signaling pathway. Our lab has shown high levels of Wnt signaling and TNC expression in anaplastic thyroid cancer (ATC), a highly lethal cancer with an abysmal ∼3-5 month median survival. Here, we investigated the role of TNC in facilitating ligand-dependent Wnt signaling in thyroid cancer. We utilized bulk RNA-sequencing from three independent multi-institutional thyroid cancer patient cohorts. TNC expression was spatially localized in patient tumors with RNA in situ hybridization. The role of TNC was investigated in vitro using Wnt reporter assays and in vivo with a NOD.PrkdcscidIl2rg-/- mouse ATC xenograft tumor model. TNC expression was associated with aggressive thyroid cancer behavior, including anaplastic histology, extrathyroidal extension, and metastasis. Spatial localization of TNC in patient tissue demonstrated a dramatic increase in expression within cancer cells along the invasive edge, adjacent to Wnt ligand-producing fibroblasts. TNC expression was also increased in areas of intravascular invasion. In vitro, TNC bound Wnt ligands and potentiated Wnt signaling. Finally, in an ATC mouse model, TNC increased Wnt signaling, tumor burden, invasion, and metastasis. Altogether, TNC potentiated ligand driven Wnt signaling and promotes cancer cell invasion and metastasis in a mouse model of thyroid cancer. Understanding the role of TNC and its interaction with Wnt ligands could lead to the development of novel biomarkers and targeted therapeutics for thyroid cancer.

## 1. Introduction

Thyroid cancer is projected to be the fourth leading cancer diagnosed in the U.S. by 2030.^1^ Most thyroid cancers are well-differentiated tumors that are treatable with surgery and radioactive iodine. However, up to 30% of patients will have metastasis, recurrence, or progression.^2,3^ Anaplastic thyroid carcinoma (ATC) is a highly lethal, aggressive form of thyroid cancer that grows quickly in the neck, leading to rapid airway compression. There has been minimal progress in treating patients with ATC, with a 12-month overall survival of less than 20% and metastasis being found at diagnosis in 50% of ATC patients.^4^ The high mortality rate of ATC is related to the rapid invasion of tumor cells into adjacent neck structures and distant metastasis of cancer cells into the lungs.^5^ An improved understanding of the mechanism of tumor cell invasion and metastasis could lead to targeted therapies and improve survival in ATC.

Technological advances, including DNA and RNA sequencing, have enhanced our understanding of many tumors, including those in the thyroid. Genomic and molecular studies have identified common driver mutations in well-differentiated thyroid cancer. ^6-18^ *BRAF*^V600E^ and *RAS* mutations are mutually exclusive and represent the most common alterations in well-differentiated thyroid cancer (WDTC).^6-18^ WDTC and ATC share driver mutations and some high-risk mutations, including *TERT* promotor and *TP53* mutations. Despite identification of high-risk mutations, clear mechanistic understanding of disease progression, invasion, and metastasis remain limited. Analysis of stromal recruitment and signaling pathway activation is needed to better understand factors that support and enhance thyroid cancer invasion and metastasis.

Wnt signaling upregulation is consistently seen in ATC across molecular subtypes.^19,20^ The Wnt/β- catenin pathway is a conserved, developmental signaling pathway that is harnessed to drive many cancers.^21-25^ Wnt signaling regulates many cellular processes, including cell fate determination, motility, polarity, and stem cell renewal.^26^ A key step in Wnt signaling is the formation of the Wnt signalosome.^25^ The Wnt signalosome is a ligand-activated receptor complex that includes the Wnt co-receptors, Frizzled and LRP6, and the cytoplasmic protein, Dishevelled.^25^ Formation of the Wnt signalosome is required for receptor-mediated stabilization of the transcriptional co-activator, β-catenin. Consequently, β-catenin accumulates in the cytoplasm, enters the nucleus, and binds the TCF/Lef1 family of transcription factors to mediate a Wnt-specific transcriptional program. Several cancers, including colorectal cancer, are known to have Wnt signaling pathway mutations that impair β-catenin degradation or mutations in proteins that promote Wnt signalosome formation.^27-29^ Other cancers are driven by increases in Wnt ligands themselves.^30-32^ Within thyroid cancer, several components of the Wnt signaling pathway are known to be altered.^19,33-36^

We recently discovered that ATCs exhibit dramatically high levels of Wnt ligand expression yet few Wnt pathway-activating mutations.^10,19^ This result suggests that Wnt signaling may play an important role in the pathophysiology of ATCs. One protein that has been shown to amplify ligand-driven Wnt signaling in development is Tenascin-C (TNC). TNC synthesis is tightly regulated in humans, having widespread expression in embryonic tissues and restricted distribution in adult tissues.^37,38^ Several studies have proposed the presence of a TNC-Wnt crosstalk during development.^39-41^ As TNC is a secreted glycoprotein, it may interact with extracellular mediators of Wnt signaling, including Wnt ligands, LRP6, or Frizzled. Studies in the whisker stem cell niche indicate that TNC can bind and concentrate Wnt-3a ligands to upregulate Wnt activation.^40^ In support of this proposed mechanism, studies of acute kidney injury demonstrated that TNC co-immunoprecipitates with Wnt-1 and Wnt-4 overexpressed in the human kidney cell line (HKC-8).^39^ TNC expression in cancer has been proposed to promote invasion and metastasis and is commonly thought to be derived from the tumor stroma.^37,41-47^ However, our understanding of TNC expression and TNC-Wnt cross-talk in thyroid cancers is limited.

Here, we investigated the expression of TNC in thyroid cancer and its role in amplifying ligand-driven Wnt signaling. In this study, we utilize patient sequencing data from three large multi-institutional thyroid patient cohorts.^9,10,48^ We demonstrate that *TNC* expression is upregulated in thyroid cancer cells along the tumor’s invasive edge and within intravascular spaces. Using co-culture and *in vitro* modeling, we demonstrate an interaction between Wnt ligand and TNC that potentiates Wnt signaling. Finally, in an ATC tumor model, we demonstrate that TNC increases Wnt pathway activation, tumor burden, tumor cell invasion, and metastasis. The interaction between TNC and Wnt is likely integral to thyroid cancer behavior and serves as a potential marker for both prognostication and targeted therapeutics.

## 2. Methods

### 2.1 Analysis of TCGA and GTEx WDTC RNA-sequencing data

GEPIA is a web-based tool that delivers rapid and customizable functions for analyzing The Cancer Genome Atlas (TCGA) and Genotype-Tissue Expression Program (GTEx) data (http://gepia.cancer-pku.cn/).^9,49^ GEPIA was used to compare TNC expression between tumor and normal samples from TCGA and GTEx. GEPIA statistical tests were performed via one-way ANOVA with a cutoff of p< 0.01. For additional analyses, TCGA Bulk RNA-sequencing and clinical data were downloaded from cBioPortal (cbioportal.org).^50-52^ The clinical data included the *BRAF*-like or *RAS*-like designation, whether the patient had lymph nodes positive for the disease at resection of the sequenced primary tumor, and the degree of extrathyroidal extension at resection (none, minimal, or moderate/advanced). TNC expression levels were log_2_ transformed and compared between TCGA tumors with *BRAF*-like versus *RAS*-like phenotypes, tumors with and without associated lymph node metastases, and tumors with no extrathyroidal extension, minimal extrathyroidal extension, or moderate/advanced extrathyroidal extension. Statistical differences in log_2_ TNC expression between groups were calculated using Wilcoxon rank-sum test with Bonferroni correction and plotted using R package ggplot2 3.5.0.^53^

### 2.2 Analysis of VUMC/UW Bulk RNA-Sequencing Patient Cohort

TNC expression was analyzed in a previously published bulk RNA-sequencing cohort of 312 thyroid resection specimens (251 patients) from Vanderbilt University Medical Center and University Washington (VUMC/UW).^10^ Samples were grouped into benign (multinodular goiters, follicular adenomas, Hürthle cell adenomas), WDTC (papillary thyroid carcinomas, follicular-variant papillary thyroid carcinomas, follicular thyroid carcinomas, Hürthle cell carcinomas), poorly differentiated thyroid carcinomas (PDTC), and ATC. WDTCs were further split into *BRAF*-like or *RAS*-like, as previously described.^9,10^ TNC expression levels were log_2_ transformed and compared across diagnoses (benign, WDTC, ATC) and between *BRAF*-like or *RAS*-like WDTCs. Within the entire malignant cohort (WDTC, PDTC, ATC) and the WDTC cohort, the expression of TNC in primary tumors was compared based on the presence or absence of associated lymph nodes and distant metastases. TNC expression was also compared between primary samples and lymph node samples. Significance was calculated using Wilcoxon rank-sum tests, and boxplots were generated with R package ggplot2 3.5.0.^53^ Progression-free survival (PFS) metrics were calculated as previously described for this cohort,^10^ using a 50^th^ percentile TNC expression cutoff within all malignant (WDTC, PDTC, and ATC) or WDTC only. R packages survival 3.5-5 and survminer 0.4.9 were used to generate PFS plots, and significance was determined via log-rank test.

### 2.3 Analysis of Lee et al. bulk RNA-sequencing patient cohort

Raw count matrices and sample meta data for 16 ATCs, 348 PTCs, and 263 normal thyroids from Lee et al. were downloaded from Gene Expression Omnibus (https://www.ncbi.nlm.nih.gov/geo/) using accession number GSE213647.^48^ Raw counts were TPM normalized and log_2_ transformed. TNC expression was compared for log2 TPM counts across diagnoses (ATC, PTC, normal thyroid) using Wilcoxon rank-sum test with Bonferroni correction and plotted using R package ggplot2 3.5.0.^53^

### 2.4 Hallmark Wnt-β-catenin score calculation

For all sequencing cohorts, hallmark Wnt-β-catenin scores were calculated from TPM (Lee, VUMC/UW) or RSEM (TCGA) counts using the R package GSVA 1.48.3 with the Molecular Signatures Database hallmark Wnt-β-catenin signaling gene set and default arguments for the GSVA function.^54,55^

### 2.5 Cancer-associated fibroblast (CAF) deconvolution

Estimated CAF levels for the VUMC/UW bulk RNA-sequencing data were previously calculated using the CAF EPIC deconvolution algorithm within TIMER 2.0.^10,56,57^

### 2.6 Bulk RNA Spearman’s correlations

Across the TCGA, Lee et al., and VUMC/UW cohorts, Spearman’s correlations between Wnt ligands, TNC, Hallmark Wnt signaling, and CAF abundance were calculated and plotted with the R packages ggplot2 3.5.0,^53^ and corrplot 0.92.

### 2.7 Multiplex immunofluorescence (IF) of formalin-fixed paraffin-embedded (FFPE) tissue

Multiplex immunofluorescence was performed as previously described.^10^ Primary antibodies (Abcam ab207178 recombinant rabbit monoclonal anti-fibroblast activation protein alpha (FAP) IgG, clone EPR20021, 1:100; Abcam ab88280 mouse monoclonal [EB2] anti-Tenascin-c 1:100; were diluted in blocking buffer and incubated on tissue sections at 4°C for 16h (Abcam, Cambridge, UK; Bioss USA, Woburn, MA). Tissue sections were washed with 0.05% Tween 20 in PBS. Secondary antibodies (Invitrogen A-21245 polyclonal goat anti-rabbit IgG Alexa Fluor 647 1:150; Abcam ab97035 polyclonal goat anti-mouse IgG H&L (Cy3 ®) preadsorbed 1:100) were diluted in blocking buffer containing Hoechst 33342 nuclear stain (1:1000) and incubated on tissue sections at 37°C for 1 h (Abcam, Cambridge, UK; Thermo Fisher, Waltham, MA). Representative 20X images were taken of each tissue section on a Nikon Spinning Disc confocal microscope.

### 2.8 RNA *in situ hybridization*

Five µm tissue sections were cut from FFPE blocks and stored at -20°C. RNAscope® probes Hs-TNC-C1, Hs-WNT2-C1, and Hs-FAP-C2 and RNAScope® 2.5 HD Duplex and RNAscope® Wash Buffer Reagents were purchased from Advanced Cell Diagnostics (Newark, CA). RNAscope® was performed according to the manufacturer’s guidelines.

### 2.9 Cell culture

K1 cells were obtained from Sigma Aldrich. WPMY-1 and HEK293 (CRL-1573) cells were obtained from American Type Culture Collection (ATCC). Cells were authenticated using STRS analysis and maintained and used experimentally at <20 passages from thaw. K1 cells were grown in RPMI (VWR) containing 10% FBS (ThermoFisher Scientific), 1% penicillin-streptomycin (Sigma), 1X MEM Non-Essential Amino Acids (VWR), and 1 mM sodium pyruvate (Vanderbilt Molecular Biology Resource). WPMY -1 and HEK293 cells were grown in high-glucose DMEM (Sigma) containing 8-10% FBS (ThermoFisher Scientific), and 1% penicillin-streptomycin (Sigma). All cell lines tested negative for Mycoplasma contamination.

### 2.10 Generating stable cell line

Stable Wnt reporter cell lines were generated using lentiviral transduction. Viral media was collected from HEK293FT cells transfected with the 7TFP lentiviral plasmid (Addgene #24308), along with the PAX2 (packaging) and pMD2G (envelope) plasmids. Thyroid cancer cell lines were cultured in lentiviral media with 8 mg/mL Polybrene for 24 hours. Antibiotic selection was performed with puromycin (10 µg/mL) (Mediatech/CellGro-Corning (MT61385RA)).

### 2.11 Transfections

Plasmid transfections were performed using Lipofectamine 3000 Transfection reagents (Invitrogen). For cell-based luciferase TOPFlash assays, K1 cells were overexpressed with either large splice variant TNC (Addgene 65414) or small splice variant TNC (Addgene 65415) and WPMY cells were overexpressed with Wnt-2 (Addgene 43809). All flag plasmids were manufactured by Gene Universal. Co-immunoprecipitation plasmid transfections were performed using the calcium phosphate method.

### 2.12 TOPFLASH assay

Cells were lysed with CellTiter-Glo 3D Assay (Promega) and One Glo Luciferase Assay (Promega). Luminescence was quantified using a Synergy NEO (BioTek multi-mode plate reader). One-Way ANOVA was performed with Tukey’s test correction.

### 2.13 Co-immunoprecipitation sample preparation

Cells were lysed using non-denaturing lysis buffer (NDLB) (50 mM Tris-HCl pH 7.4, 300 mM NaCl, 5 mM EDTA, and 1% Triton X-100 (w/v), supplemented with 1 mM PMSF and PhosSTOP phosphatase inhibitor cocktail tablets(Roche). Samples were incubatedat 4°C for 30 min, followed by clarification by spinning in a microfuge at 13,000 RPM for 10 min at 4°C. Lysates were diluted to 1 mg/mL with NDLB and incubated with antibodies with overnight rotation at 4°C. Samples were then incubated with Protein A/G magnetic beads (Millipore) for two hours with rotation at 4°C. Beads were washed 5 times with NDLB, and sample buffer was added to elute the bound protein (95°C for 10 min). Proteins were analyzed by SDS-PAGE and immunoblotting. Fluorescence signal was detected using a an Odyssey (LI-COR). Obtained images and band intensity were analyzed using Empiria (LI-COR).

### 2.14 Co-Immunoprecipitation sample preparation for recombinant proteins

Recombinant TNC (1 µg, EMD Millipore) and Recombinant Wnt-2 (Biomatik) were diluted to a total volume of 1 mL with NDLB and incubated with TNC antibody overnight at 4°C. Samples were then incubated with Protein A/G magnetic beads (Millipore). Beads were washed 5 times with NDLB, and sample buffer was added to elute the bound protein (20°C for 1h). Proteins were analyzed by SDS-PAGE and immunoblotting. Chemiluminescence signal was detected using an Odyssey (LI-COR). Obtained images and band intensity were analyzed using Image Studio (LI-COR).

### 2.15 Antibodies for immunoblotting

The following antibodies were used for immunoblotting: Rabbit anti-FLAG (Proteintech, 20543-1-AP), Mouse anti-FLAG (Vanderbilt Protein and Antibody Resource), Mouse anti-Tubulin (Developmental Studies Hybridoma Bank, E7), Mouse anti-Tenascin C (Abcam, ab3970), Rat anti-Tenascin C (R&D Systems, MAB2138-SP), Rabbit Anti-Wnt-2 (Abcam, ab1009222), Rabbit anti-V5 (Cell Signaling, 13202S), Goat anti-rat IgG H + L-HRP (Thermo Fisher, 31470), Goat anti-mouse IgG H + L-HRP (Promega, W4021), Goat anti-rabbit IgG H + L-HRP (Promega, W4011), Goat anti-rabbit 800 (Licor), Donkey anti-mouse 800 (Licor), Goat anti-rabbit 680 (Licor), and Goat anti-mouse 680 (Licor). All primary antibodies were used at 1:1000 dilution except anti-tubulin (1:10000) and anti-flag (1:2000). All secondary HRP antibodies were used at 1:2000 dilution, and all fluorescence antibodies were used at 1:20000 dilution.

### 2.16 Mouse experiments

All procedures were approved by the Institutional Animal Care and Use Committee prior to completion. NOD.PrkdcscidIl2rg-/- (NSG-Jackson Laboratories) were injected with 1×10^6^ xenograft THJ-16T cells subcutaneously in the flank using a 25G SubQ needle affixed to a 1 mL syringe. When tumors became palpable (approximately 1-week post-injection), intratumoral injections were performed with 1X PBS (Corning) or recombinant Tenascin-C (EMD Millipore, 0.2 mg/mL) twice weekly. Tumors were measured twice weekly using digital calipers, and mice were weighed weekly to ensure that weight loss did not exceed 20% of body weight. When tumors reached 2 cm in any dimension or ulcerated, mice were humanely euthanized. Both male (14 animals) and female (11 animals) mice were used in all experiments and was not considred a factor in statistical analysis.

### 2.17 Mouse tumor histology and immunohistochemistry

Tumors were resected and weighed, fixed in 10% neutral buffered formalin, processed, and embedded in paraffin. 5 µm sections were cut and either stained with hematoxylin and eosin or used for immunohistochemistry. Slides used for immunohistochemistry were stained by the Translational Shared Pathology Resource (TPSR) center. Slides were placed on the Leica Bond Max IHC stainer. Heat induced antigen retrieval was performed using Epitope Retrieval 2 solution for 20 minutes. Slides were placed in a Protein Block (Ref# x0909, DAKO) for 10 minutes. Slides were incubated with anti-B-Catenin (Cell Signaling, 9582) for one hour at a 1:100 dilution. The Bond Polymer Refine detection system was used for visualization. Slides were the dehydrated, cleared and coverslipped.

### 2.18 Statistics

Tumor volumes and tumor weights from three experiments (batches) were analyzed. Linear mized-effects model was used to evaluate the association between treatment and Wnt reporter activation and account for correction due to technical replicates from the same biological samples. Wnt reporter activation by treatment group was estimated using least-squares means (i.e., model-based means), and differences among groups were compared using the Wald test with Tukey’s test correction. We employed the power variance function to address heteroscedasticity across treatment groups. Batch was included as a covariate to adjust for potential batch effects. Residual analysis was conducted to verify the model’s assumptions. Tumor volumes were log-transformed (with the addition of 1 to avoid log(0)) to address heterogeneity detected in residual analysis. Statistical analyses were performed in R v4.4.0.

## 3. Results

### 3.1 TNC expression is increased in WDTC and ATC

To first analyze Tenascin-C expression in thyroid cancer, we assessed TCGA WDTC (n = 512) and GTEx normal thyroid (n = 337) gene expression using GEPIA. *TNC* expression was upregulated in WDTC compared to normal thyroid tissue (**Fig. 1A,** p<0.01). Additionally, *TNC* is known to have several different splicing events,^58^ so we evaluated the different isoforms of TNC present in thyroid cancer. TNC isoforms-12, 11, and 1 are the most common (**Fig. S1A** and **Fig. S1B**). We next investigated *TNC* expression in the more aggressive thyroid cancer, ATC. Using publicly available sequencing data from Lee et al., we assessed *TNC* expression in normal, PTC (the most common type of WDTC), and ATC samples. We found that *TNC* expression is increased in PTC relative to normal (p<0.001) and ATC relative to normal (p<0.001), with the highest expression in ATC (**Fig. 1B**).

**Figure 1.**
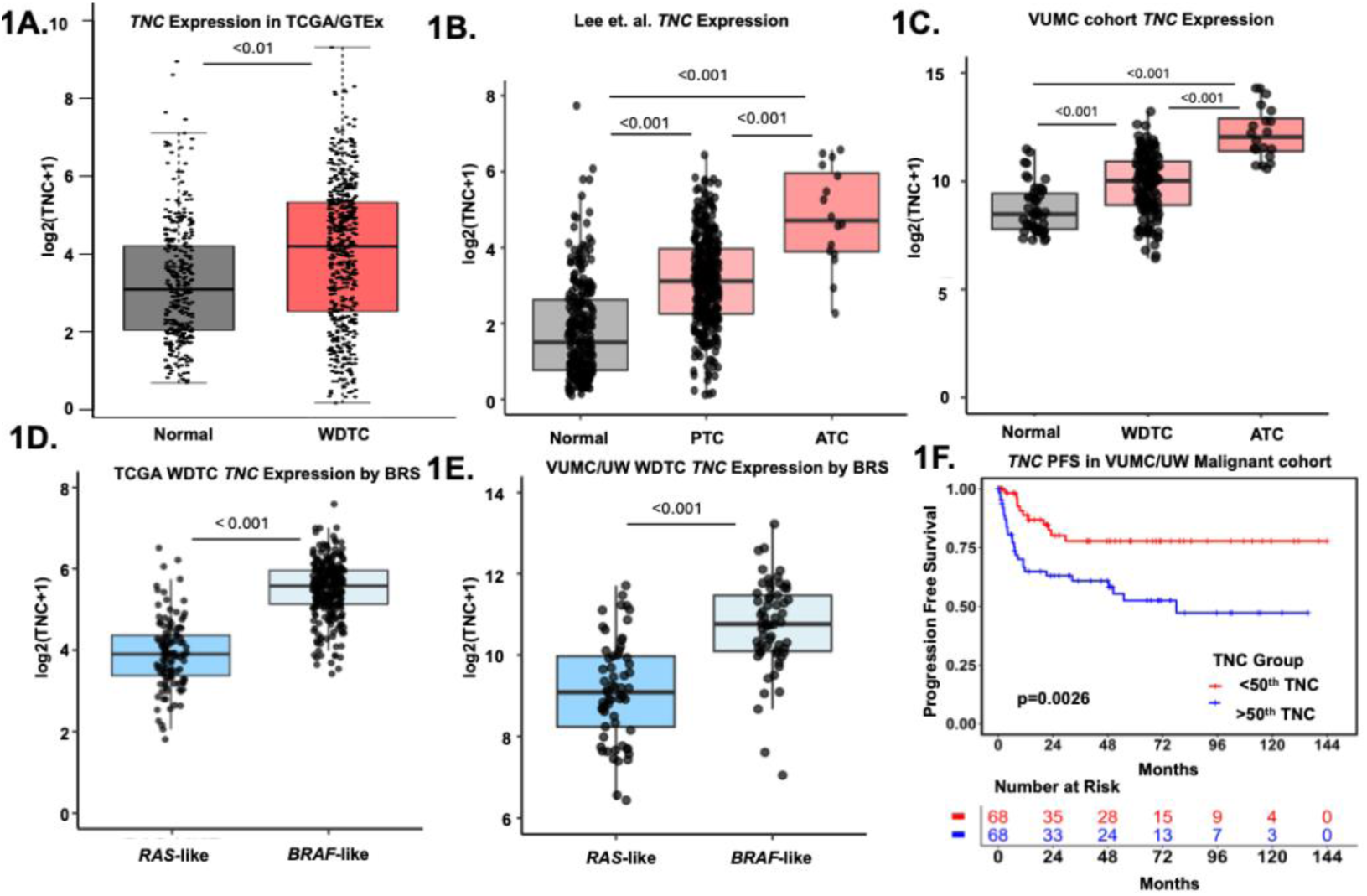
TNC expression is higher in thyroid cancer patient samples. **(A)** Boxplots generated with GEPIA showing TNC expression in TCGA and GTEx thyroid samples split by normal thyroid versus WDTC. **(B-C)** Boxplots showing *TNC* expression in **(B)** Lee et. al. showing normal, PTC, and ATC samples and **(C)** primary samples by diagnosis (benign = n of 46; well-differentiated thyroid cancer (WDTC) = n of 136; anaplastic thyroid cancer (ATC) = n of 20. p-values calculated using using Wilcoxon rank-sum test with Bonferroni correction. **(D-E)** *TNC* expression based on *BRAF*-like or *RAS*-like gene expression phenotypes in **(D)** TCGA WDTC Cell 2014 cohort and **(E)** VUMC/UW primary WDTCs. p-values calculated using using Wilcoxon rank-sum test with Bonferroni correction. **(F)** Progression-free survival (PFS) plot for all malignant patients with less than 50^th^ percentile *TNC* expression (red) or greater than 50^th^ percentile TNC expression (blue) in primary tumors. p-values calculated using log-rank test.

We recently collected and published an analysis of a large cohort of whole exome and bulk RNA-sequencing data analysis of benign and malignant thyroid tissue.^10^ This cohort was enriched for ATC and WDTCs with metastases and poor outcomes to improve detection of drivers of aggressive disease. Similar to the cohort from Lee et. al., TNC expression is increased in WDTC (p<0.001) and ATC samples (p<0.001), with the highest increase in ATC (**Fig. 1C**). We further classified WDTCs into *RAS*-like or *BRAF*-like based on their gene expression profiles.^9,10^ Using the TCGA WDTC cohort, *TNC* expression is increased in *BRAF*-like WDTCs compared to *RAS*-like tumors (**Fig. 1D**, p<0.001). Similar to TCGA WDTCs, we observe a significant increase in our cohort in *TNC* expression within WDTCs that are *BRAF*-like compared to *RAS*-like (**Fig. 1E**, p<0.001). Taken together, *TNC* expression is upregulated in thyroid cancer compared to normal tissue. The highest increases in TNC are observed in *BRAF*-like WDTCs and ATCs. Finally, using PFS analysis of our whole malignant cohort, we see an improved survival in patients with low *TNC* expression (split by 50^th^ percentile, **Fig. 1F**, p=0.0026). This significance is not seen within our WDTC cohort (**Fig. S1C**, p=0.46), suggesting that this survival difference is likely driven, at least in large part, by the high TNC expression in lethal ATCs.

### 3.2 TNC expression is increased in metastasis

TNC expression has been implicated as a driver of metastasis,^59^ so we further assessed the association between *TNC* and metastasis in our thyroid cancer cohorts. Within the TCGA WDTC cohort^9^, primary tumors from patients with lymph node metastases showed higher overall *TNC* expression compared to primary tumors from patients without lymph node metastases (**Fig. 2A**, p=0.0026). Additionally, *TNC* expression was increased in tumors with minimal (p<0.001) and moderate/advanced (p<0.001) extrathyroidal extension (**Fig. 2B**). An evaluation of TNC and metastasis within the VUMC/UW cohort shows similar findings. *TNC* expression is increased in the primary tumors of patients with lymph node metastases (**Fig. 2C,** p<0.001) when evaluating our whole malignant cohort as well as when we restricted the cohort to only WDTC (**Fig. S2B,** p<0.001). This analysis confirms that our correlation is not driven solely by ATC. While TNC expression in the primary tumors also correlates with distant metastatic disease, this trend is not significant (**Fig. S2A**, **S2C**).

**Figure 2.**
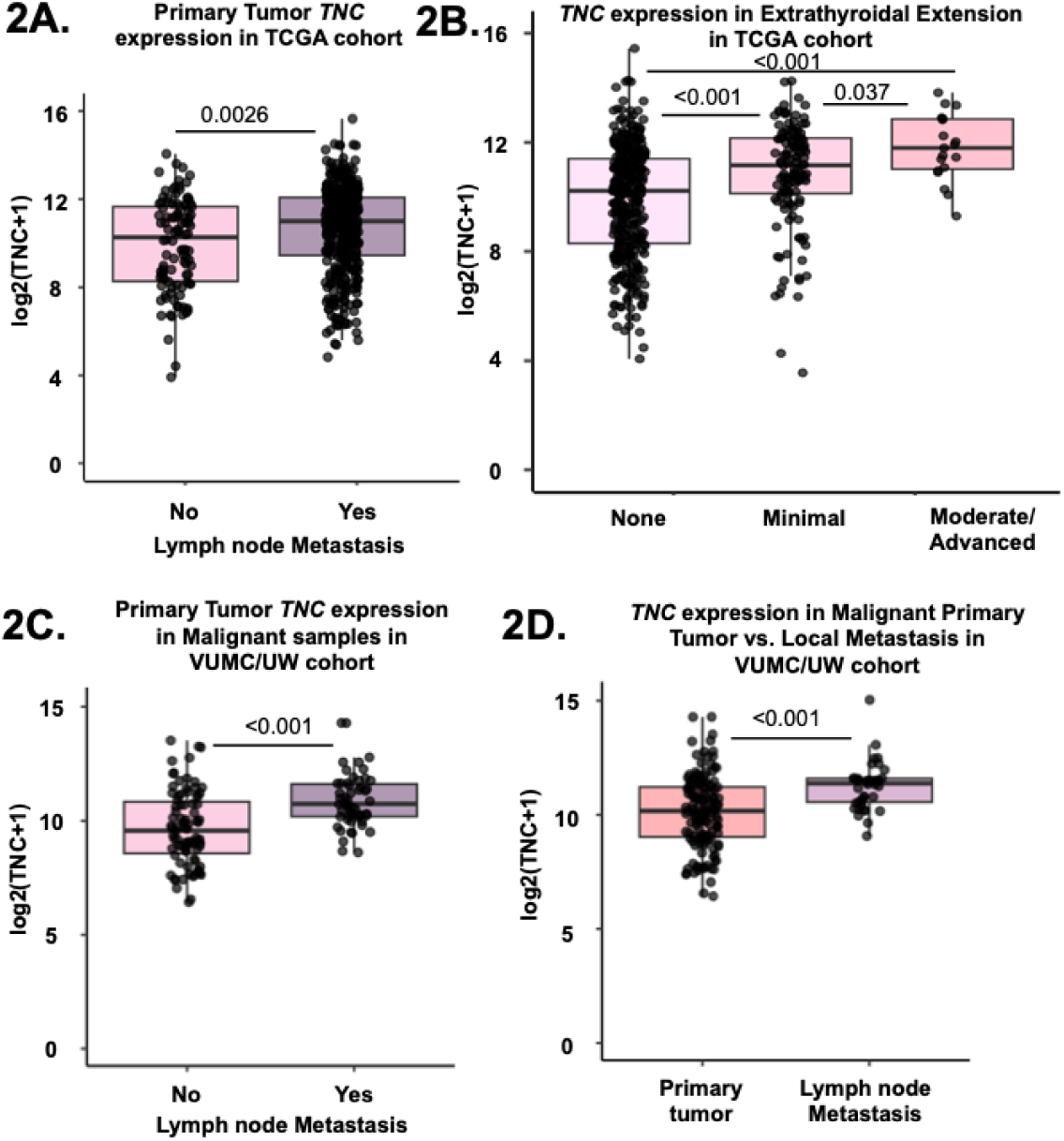
TNC expression is correlated with local invasion and lymph node metastasis. **(A-B)** Boxplots showing TNC expression in TCGA WDTCs split by **(A)** presence (YES) or absence (NO) of lymph node spread and **(B)** extent of extrathyroidal extension (none, minimal, moderate/advanced). p-values calculated using Wilcoxon rank-sum test with Bonferroni correction. **(C)** Boxplots showing TNC expression in primary malignant tumors from the VUMC/UW cohort split by presence (YES) or absence (NO) of associated local metastases. **(D)** Boxplots showing TNC expression in primary malignant tumors versus lymph node metastases in the VUMC/UW cohort. p-values calculated using Wilcoxon rank-sum test with Bonferroni correction.

Interestingly, not only is TNC increased within the primary tumors of patients with lymph node metastases, but it is also increased within the metastatic cells of the lymph nodes (**Fig. 2E,** p<0.001 and **Fig. S2D**, p<0.001).

In conclusion, across multiple published thyroid cancer sequencing cohorts, we observe an increase in TNC in malignant samples. This increase is highest in samples with aggressive behavior, including lymph node metastases, extrathyroidal extension, and transformation to ATC.

### 3.3 Spatial localization of RNA and protein expression of TNC in ATC

While it is commonly thought that TNC is expressed by fibroblasts in the tumor microenvironment,^44,45^ recent studies also suggest that tumor cells in both breast and head and neck cancer can produce TNC.^46,47^ To identify which cells are making TNC in thyroid cancer and their spatial localization, we performed RNA *in situ* hybridization and multiplex immunofluorescence on patient ATCs. First, using RNA *in situ* hybridization, we probed for *TNC* and fibroblast activation protein alpha (*FAP*), a ubiquitous CAF marker. We found that 7 of 12 samples (58%) have *TNC* staining of tumor cells along the invasive edge **(Fig. 3A** and **3B**). In addition, 10 of 12 samples (83%) have *TNC* staining *of* endothelial cells and tumor cells within blood vessels (intravascular invasion, **Fig. 3C** and **3D**). Finally, all 12 samples (100%) showed fibroblast TNC staining **(Fig. 3A** and **3B)**. Confirmation of protein expression using multiplex immunofluorescence staining shows TNC protein expression of invading tumor cells along the tumor-stromal border (**Fig. 3E**). Altogether, we observe TNC expression by tumor cells at the tumor-stromal interface in areas of stromal and vascular invasion. We also detect TNC expression within the fibroblast-rich stroma.

**Figure 3.**
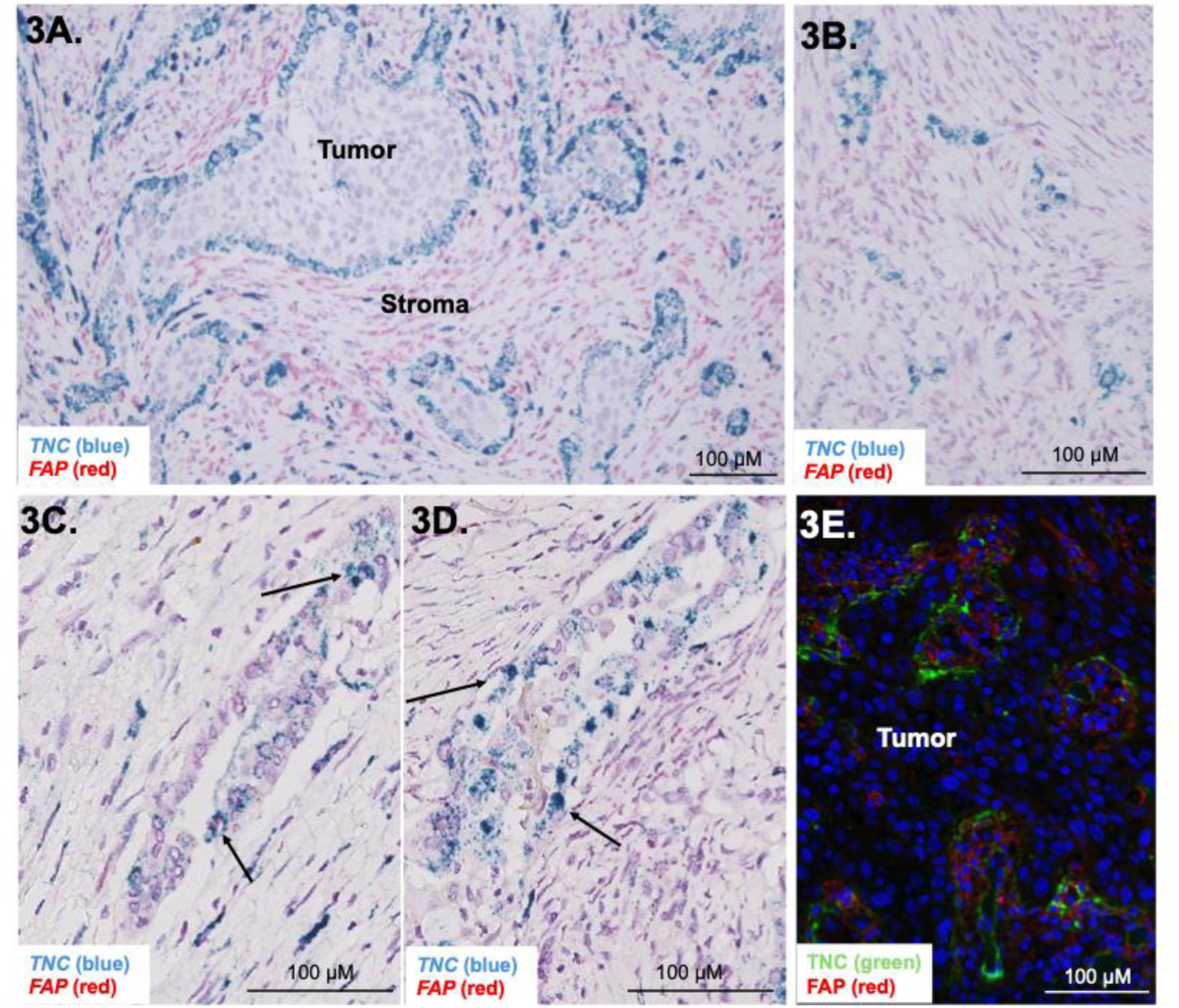
TNC expression is along the epithelial stromal border in ATC. **(A-B)** RNAscope® of representative ATC samples showing staining for TNC (blue) or Fibroblast activating protein (FAP, red). **(C-D)** RNAscope® of representative PTC regions within ATC samples showing staining for TNC (blue) or FAP (red). **(E)** Multiplex-immunofluorescence staining of representative ATC sample showing TNC staining (green), FAP (red), and Hoechst nuclear stain (blue).

### 3.4 TNC and Wnt-2 expression correlate in thyroid cancer

Although TNC is found at the invasive border of ATC, the mechanism by which it influences tumor invasion is unclear.^46,47^ In non-neoplastic disease, TNC has been shown to interact with Wnt ligands and augment ligand-dependent Wnt signaling.^39,41^ As aggressive thyroid tumors have increased expression of Wnt ligands, we hypothesized that TNC acts to enhance Wnt signaling at the invasive border of thyroid tumors through interactions with Wnt ligands. To investigate the relationship between TNC and Wnt signaling, we first looked at hallmark Wnt/β-catenin signaling gene activity and observed a positive correlation with *TNC* expression in all malignant samples using the Lee et. al. cohort (**Fig. 4A**, p<0.001) and our VUMC/UW cohort (**Fig. 4B**, p<0.001). We observed the same positive correlation when we looked at WDTC *TNC* expression and Hallmark Wnt/β-catenin signaling gene activity across all patient cohorts (**Fig. S3A-C**).

**Figure 4.**
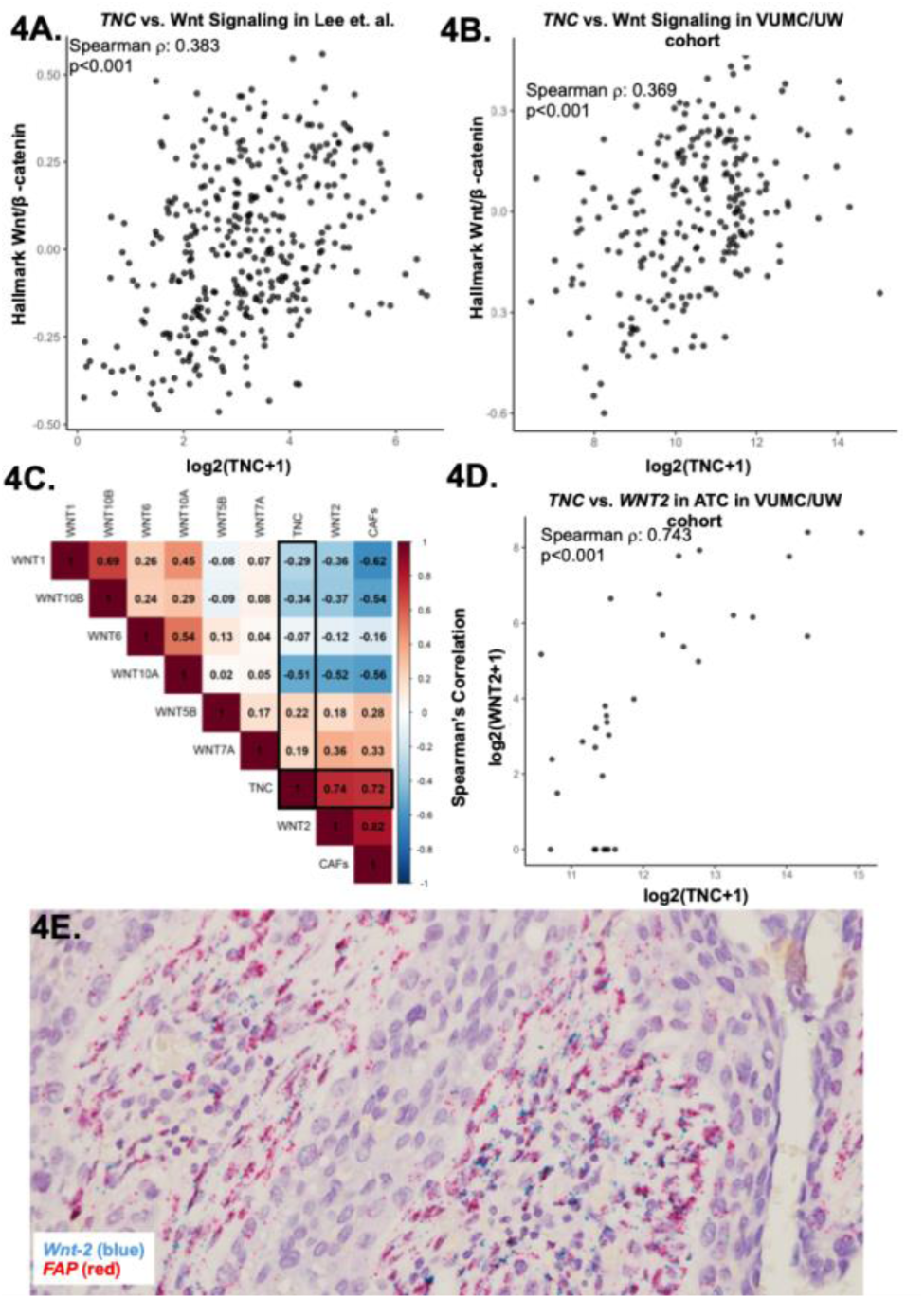
TNC and Wnt-2 expression are correlated in ATC. **(A-B)** Spearman’s correlation between hallmark Wnt/β-catenin gene activity in **(A)** Lee et. al. or **(B)** VUMC/UW bulk RNA-sequencing cohorts for all malignant samples. **(C)** Plot of Spearman’s correlations between Wnt ligands, Cancer-associated fibroblasts (CAFs), and TNC in VUMC/UW ATCs. **(D)** Spearman’s correlation between TNC and *WNT2* in VUMC/UW ATC samples. **(E)** RNAscope® of representative ATC sample showing staining for *WNT2* (blue) or *FAP* (red).

Next, we aimed to determine if there are specific Wnt ligands that TNC is interacting with to potentiate Wnt signaling. We recently identified seven Wnt ligands (Wnt-1, Wnt-2, Wnt-5b, Wnt-6, Wnt-7a, Wnt-10a, Wnt-10b) that are upregulated in ATC.^19^ Because we spatially observed TNC next to fibroblasts, we chose to look at the correlation between TNC, ATC-upregulated Wnt ligands, and CAF abundance in ATCs (**Fig. 4C**). We identified a markedly stronger correlation between *TNC* and *WNT2* than any other Wnt ligand (**Fig. 4C-D**, ρ=0.74, p<0.001). Additionally, both *TNC* and *WNT2* were correlated with CAFs (ρ=0.72 and 0.82, respectively). We confirmed the association between *TNC* and *WNT2* across malignant and WDTC-restricted cohorts from Lee et al., TCGA, and VUMC/UW (**Fig. S3D-H**). As multiple reports have indicated that Wnt-2 is expressed by CAFs, we anticipated that Wnt-2 may be interacting with TNC via CAFs adjacent to invasive, TNC-expressing tumor cells.^60-62^ To identify whether Wnt-2 is produced in close proximity to the invasive border of tumor cells, we performed RNA *in situ hybridization*, probing for *WNT2* and *FAP* within ATC tumors. We found that *WNT2* is made by CAFs within the tumor microenvironment adjacent to the leading edge of tumor cells (**Fig. 4E**). In conclusion, across patient cohorts, TNC is correlated with hallmark Wnt/β-catenin gene activity. Amongst Wnt ligands, TNC exhibits the strongest correlation with Wnt-2, which is produced by CAFs at the invasive border with tumor cells, suggesting a possible interaction between Wnt-2 and TNC.

### 3.5 TNC potentiates Wnt signaling and directly interacts with Wnt-2

TNC has been shown to influence Wnt signaling by interacting directly with Wnt ligands.^39,41^ To study the crosstalk between TNC and Wnt ligands *in vitro*, we used a TOPFLASH K1 thyroid cancer cell line. First, we tested whether TNC could potentiate ligand-driven Wnt signaling using a co-culture method. Specifically, we overexpressed TNC in the TOPFLASH K1 thyroid cancer cell line and overexpressed Wnt-2 in a fibroblast cell line, WPMY. With no overexpression, or with only TNC overexpression, we observe baseline activation of the Wnt signaling pathway. With Wnt-2 overexpression, we see activation of Wnt signaling. However, Wnt signaling is potentiated when we overexpress both TNC and Wnt-2, independent of whether the small or large TNC splice variant is overexpressed (**Fig. 5A-B**). These data suggest a role for TNC in potentiating Wnt expression in the tumor microenvironment through Wnt-ligand mediated activation.

**Figure 5.**
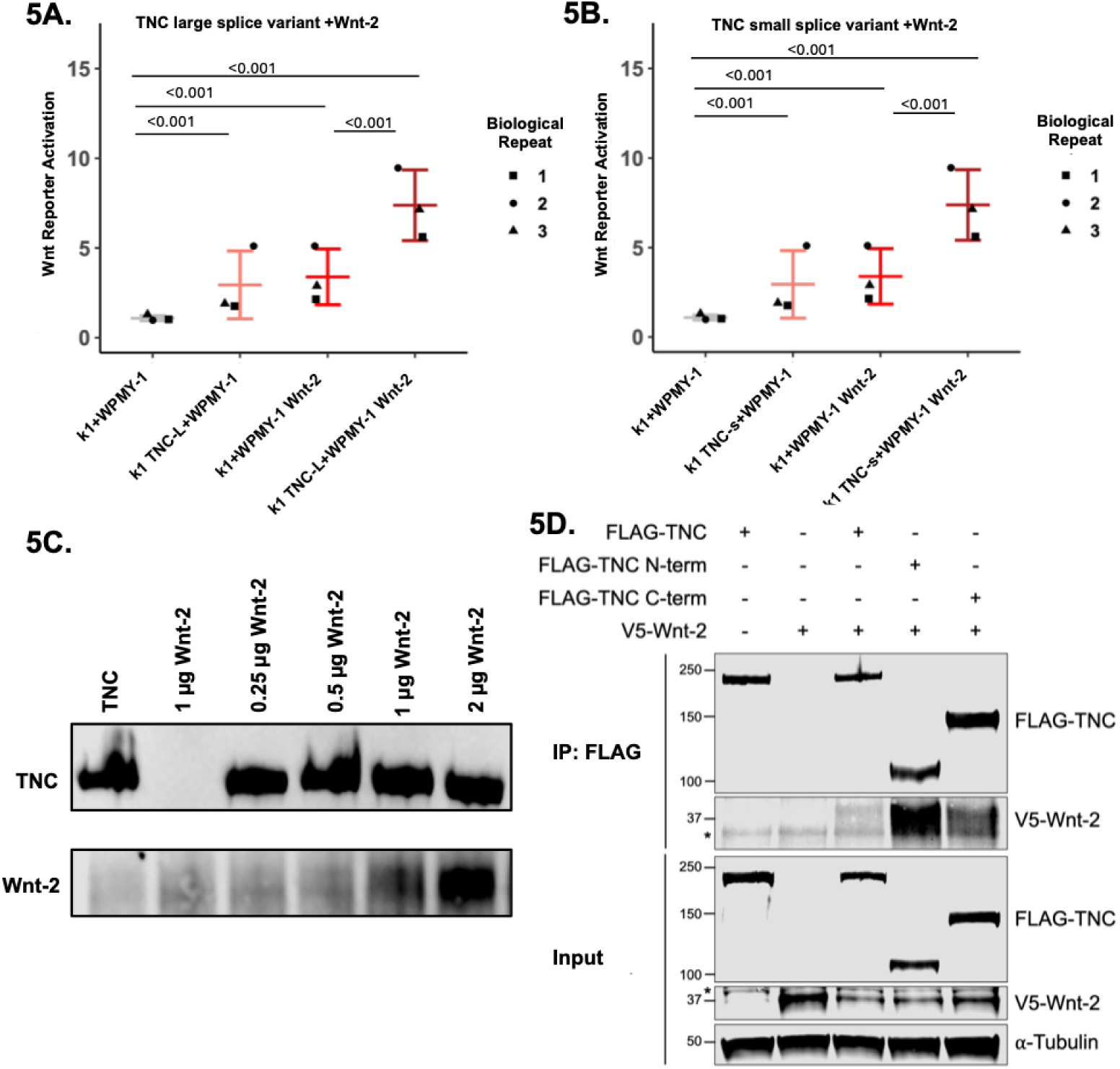
TNC variants potentiate Wnt signaling in presence of Wnt ligands. **(A-B)** TOPFLASH Wnt reporter readouts for K1 cells overexpressing **(A)** TNC large splice variant or **(B)** TNC small splice variant co-cultured with Wnt-2 expressing WPMYs. p-values were calculated using Wald test with Tukey’s test correction. **(C)** TNC and Wnt-2 co-immunoprecipitation using several concentrations of recombinant Wnt-2 and 0.1 mg/mL of recombinant TNC. **(D)** Flag-tagged TNC and V5-tagged Wnt-2 co-overexpressed in HEK293 cells was immunoprecipitated (IP) with anti-FLAG antibody and protein A/G beads, and co-immunoprecipitated V5-Wnt2 was detected by immunoblotting. Abbreviations: WCL, whole cell lysates. * indicates non-specific band

To further explore an interaction between TNC and Wnt ligand, we used recombinant TNC and recombinant Wnt-2 and performed co-immunoprecipitation experiments, in which we pulled down TNC and immunoblotted for Wnt-2. Wnt-1, 3a, and 4 have previously been shown to co-immunoprecipitate with TNC, but the interaction between TNC and Wnt-2 has not been previously described.^39,40^ We found that Wnt-2 pulled down with recombinant TNC, indicating a potential interaction between the two proteins (**Fig. 5C**). To explore the interaction between TNC and Wnt-2 further, we FLAG tagged the small splice variant of TNC and created two distinct TNC truncation constructs consisting of either the N-terminus, containing EGF-like repeats, or the C-terminus, containing FN3-like repeats (**Fig. S1B**). We then co-overexpressed each constructof TNC with V5-tagged Wnt-2and performed co-immunoprecipitation experiments. We found that Wnt-2 immunoprecipitated with all three TNC constructs, providing further support for the interaction between TNC and Wnt-2. Wnt-2 was pulled down most strongly with the N-terminus of the small splice variant of TNC, indicating that the Wnt-2 binding site on TNC is likely contained within the N-terminal region (**Fig. 5D**). Surprisingly, much weaker pulldown was observed with the full-length TNC fragment, suggesting that the binding sites for Wnt ligands on TNC may be partially conformationally masked.

### 3.6 TNC increases tumor burden *in vivo*

Finally, we explored the role of TNC in thyroid tumor growth and invasion *in vivo*. To do so, we injected the THJ-16T patient-derived ATC xenograft cell line subcutaneously into the flanks of NSG mice. THJ-16T forms a similar morphology tumor to the ATC we commonly observe in TNC-positive patient tumors, with nests of squamoid tumor cells surrounded by CAFs (**Fig. S4A-B**). After the tumors became palpable, ∼seven days after injection, either PBS or recombinant TNC was injected twice weekly into the tumors of these mice, and tumor size was measured. We saw a significant increase in tumor size (p=0.008) and weight (p=0.003) in our TNC mice compared to our PBS controls (**Fig. 6A** and **6B**). In addition, in 4 TNC-treated mice, we observed dramatic cancer cell invasion that extended into the ribcage, pleura, lung, peritoneum, and liver. Metastatic disease was also present in 3 TNC-treated mice with tumors present in the spleen and lymph nodes (**Fig. 6C-D, S4C**). PBS-treated tumors were only found subcutaneously in the skin of injected mice. After the humane endpoint was reached, tumors were collected for histologic evaluation and β-catenin immunohistochemistry. Immunohistochemical staining demonstrated membranous β-catenin staining with rare cytoplasmic staining in PBS-treated tumors (**Fig. 6E**). However, strong nuclear and cytoplasmic β-catenin staining was identified in mice treated with recombinant TNC, indicating activation of the Wnt pathway in TNC-treated tumors (**Fig. 6F**). Taken together, these findings show that TNC treatment led to an increase in Wnt activation, tumor burden, tumor cell invasion, and tumor metastasis.

**Figure 6.**
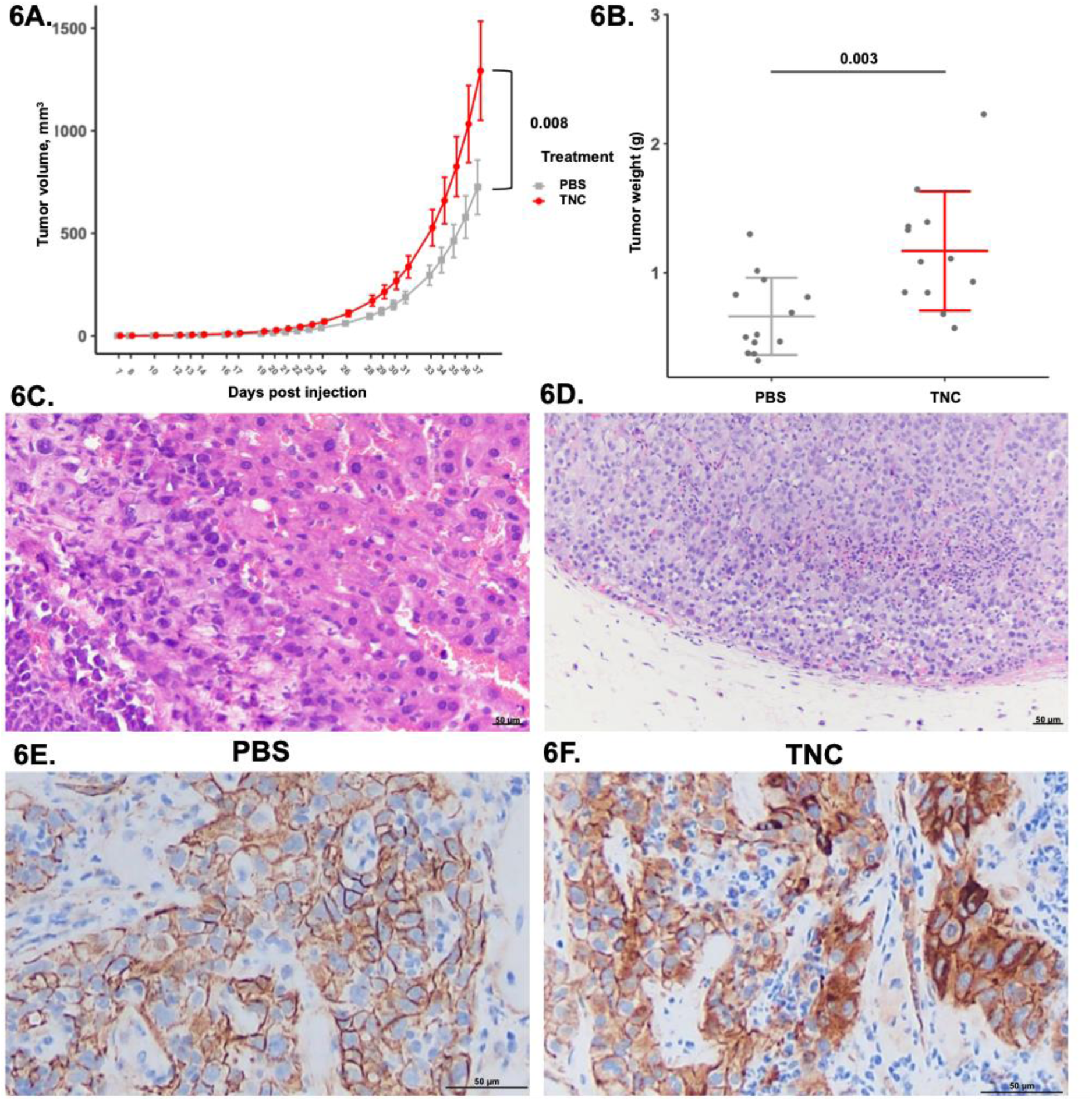
TNC expression increases tumor burden *in vivo*. **(A)** THJ-16T tumor burden (mm^3^) measured over time for tumors with PBS injections into tumor versus tumors with recombinant TNC injections into tumor. p-values calculated using linear mixed -effect model. **(B)** THJ-16T tumor weight after resection. p-values calculated using one way ANOVA. **(C)** H&E of THJ-16T tumor invading the liver of TNC-treated mouse. **(D)** H&E of lymph node from TNC-treated mouse that is completely replaced by 16T-PDX tumor. **(E)** Immunohistochemistry of β-catenin from PBS-treated THJ-16T tumors. **(F)** Immunohistochemistry of β-catenin from TNC-treated 16T-PDX tumors.

## 4. Discussion

TNC is expressed widely in development but is largely silenced in adulthood. In this study, using multiple large patient cohorts, *in vitro* models, and *in vivo* mouse models, we identified *TNC* as an important modulator of Wnt signaling in thyroid cancer. Our data demonstrate that TNC expression is tightly regulated in thyroid cancer, as it is expressed by tumor cells along the leading edge and within areas of intravascular invasion. The TNC-expressing tumor cells along the leading edge are immediately adjacent to Wnt-2 producing fibroblasts. This close approximation allows for a TNC-Wnt interaction that leads to the potentiation of Wnt signaling within both the invading tumor cells and adjacent stromal cells. Based on our *in vivo* animal studies, we speculate that this potentiation of Wnt signaling by TNC plays an important role in driving thyroid cancer invasion and metastasis. Until now, few drivers of thyroid cancer invasion have been identified. Thus, TNC may then serve as an important marker that could be used to inform both prognosis and treatment.

This study also has broad implications beyond thyroid cancer. TNC had been previously shown to interact with Wnt ligands in a handful of developmental or fibrosis models. We show here that within thyroid cancer there is an interaction between *TNC* and *Wnt-2* that leads to pathway activation. To our knowledge, we are the first to map the interaction between TNC and Wnt-2. The existence of a new Wnt modulator could lead to new research directions for cancer therapeutics and beyond. First, TNC is expressed in many tissues and is important in metastasis and several pro-tumorigenic roles. Specifically, in breast cancer, TNC splice variants have been known to increase invasion.^63^ Further, the mechanism behind TNC’s pro-oncogenic effect may be through its interaction with Wnt ligands. Within pancreatic cancer, a crosstalk between TNC and Wnt signaling has already been established.^64^ The ability of TNC to modulate Wnt signaling across a variety of cancers could lead to the development of target therapies for both TNC and Wnt. Second, Wnt-2 is expressed in many tissues and is important in development and tissue patterning. In the liver, Wnt-2 is expressed by endothelial cells in the central vein and is essential for regulating liver zonation.^65^ TNC expression has also been identified in inflammatory models of liver disease, fatty liver disease, fibrotic liver disease, and liver transplant rejection.^66,67^ The ability of TNC to modulate Wnt signaling across a variety of human diseases could lead to new therapeutic directions for many Wnt-driven disease processes.

There are several important limitations of our study. First, analysis of TNC levels in tumor cohorts is performed using bulk sequencing. Such a sequencing approach limits the ability to detect the specific cell types that express TNC and Wnt-2. To overcome this limitation, we have performed RNAScope and multiplex immunofluorescence to localize their expression within patient tissues. Another limitation of our study is that both of our TNC variants (N-terminus and C-terminus) include the fibrinogen-like globe domain. As both variants pull down Wnt-2, it is possible that the fibrogen-like globe domain binds Wnt-2. Additional studies are needed to precisely elucidate the exact site of Wnt-2-TNC interaction. Finally, our *in vivo* mouse studies utilize a flank model of ATC. This model was chosen over an orthotopic model since neck ATCs lead to rapid airway compression, limiting our ability to perform studies on tumor growth and metastasis. However, a limitation of the flank model is that the tumor vasculature may not produce the same metastatic disease as an orthotopic model. Despite this limitation, TNC-treated mice with flank tumors showed metastasis to draining lymph nodes and spleen.

## 5. Conclusion

Tenascin-c, a hexameric glycoprotein, has been known to increase tumor invasion and metastasis in non-thyroid cancers. Using multiple patient cohorts with RNA sequencing data, we show that thyroid cancer expresses elevated levels of tenascin-C. Within local metastasis, ATCs, and *BRAF*-mutant WDTCs, we observe increased *TNC* expression. Using RNA *in situ hybridization* and multiplex immunofluorescence, we are able to localize this expression within the tissue. We see a striking pattern emerge with *TNC* expression in a single-cell layer along the epithelial-stromal border and expressed by tumor cells themselves. Additional TNC expression is identified at sites of intravascular invasion, directly linking TNC expression with metastasis. Mechanistically, we show that both the large and small splice variants of TNC bind to Wnt-2 ligand and potentiate Wnt signaling. Finally, *in vivo* mouse tumors show an increase in tumor size, tumor invasion, metastasis and Wnt activation when recombinant TNC is injected intratumorally. Taken together, we propose that TNC potentiates Wnt signaling and increases thyroid cancer tumor burden. These findings suggest that TNC may be a useful biomarker of thyroid cancer invasion and metastasis. In addition, targeting TNC may lead to therapeutic advances for some of the most aggressive and lethal thyroid cancers.

## Ethical Considerations

### Declaration of Interests

E.L. is a co-founder of StemSynergy Therapeutics, a company that seeks to develop inhibitors of major signaling pathways (including the Wnt pathway) for the treatment of cancer.

### Funding

This research was funded by NIH T32ES007028 (H.A.H.); NCI 1F30CA281125-01 (M.A.L.); NIH T32GM007347 (M.A.L.); NIH T32 T32GM008554 (A.S); NIH R35GM122516, R01CA244188,R01CA272875 (E.L.); VCORCDP K12CA090625, K08CA240901, and R01CA272875 (V.L.W.); ATA 2019-0000000090 (V.L.W.); ACS RSG-22-084-01-MM (V.L.W.); and ACS 133934-CSDG-19-216-01-TBG (V.L.W.). The publication described was additionally supported by CTSA award No. UL1 TR002243 from the National Center for Advancing Translational Sciences. Its contents are solely the responsibility of the authors and do not necessarily represent official views of the National Center for Advancing Translational Sciences or the National Institutes of Health.

## Acknowledgements

The authors would like to thank Dr. John Copland for providing THJ-16T cells for these studies. We thank the Vanderbilt Translational Pathology Shared Resource (TPSR) for their support in tissue sectioning and staining (NCI/NIH Cancer Center Support Grant P30CA068485 and the Shared Instrumentation Grant S10 OD023475-01A1 for the Leica Bond RX), the Vanderbilt Technologies for Advanced Genomics (VANTAGE) core facility for their support for sequencing (5P30 CA68485-19, S10 OD023475-01A1, DK20593, DK58404, DK59637, and EY08126), and the Vanderbilt Cell Imaging Shared Resource (CISR) for their support with confocal microscopy (supported by NIH grants CA68485, DK20593, DK58404, DK59637, EY08126, and S10MH130456-01). Figure S1B was created using BioRender.

## Author Contributions

The conception and design of the study were performed by V.L.W. and E.L. The manuscript was written by H.A.H. and M.A.L. Material preparation and data collection were performed by H.A.H., M.A.L., G.J.X., A.C.S., C.J.P., K.C., H.C., A.H., D.D., M.T., J.G., S.M.S., Q.S., J.L.N., S.L.R., C.C.S., L.A.B., N.B., B.A.M., J.H.C., E.C.H., and P.H. Sequencing analysis was performed by H.C., Q.S., M.A.L., and G.J.X. Statistical analysis was performed by F.Y. and S.C.C. All authors read and approved the final report.

Code for all analyses is available at the following link on GitHub: https://github.com/xgj797/Molecular-Signature-Incorporating-Immune-Microenvironment-Enhances-Thyroid-Cancer-Outcome-Prediction, and https://github.com/hartheat/Tenascin-c-potentiates-Wnt-signaling-in-thyroid-cancer. Any additional information required to reanalyze the data reported in this paper is available from the lead contact upon request.

## Notes

https://github.com/hartheat/Tenascin-c-potentiates-Wnt-signaling-in-thyroid-cancer

