## Supplemental Figures for "Tenascin-C potentiates Wnt signaling in thyroid cancer"

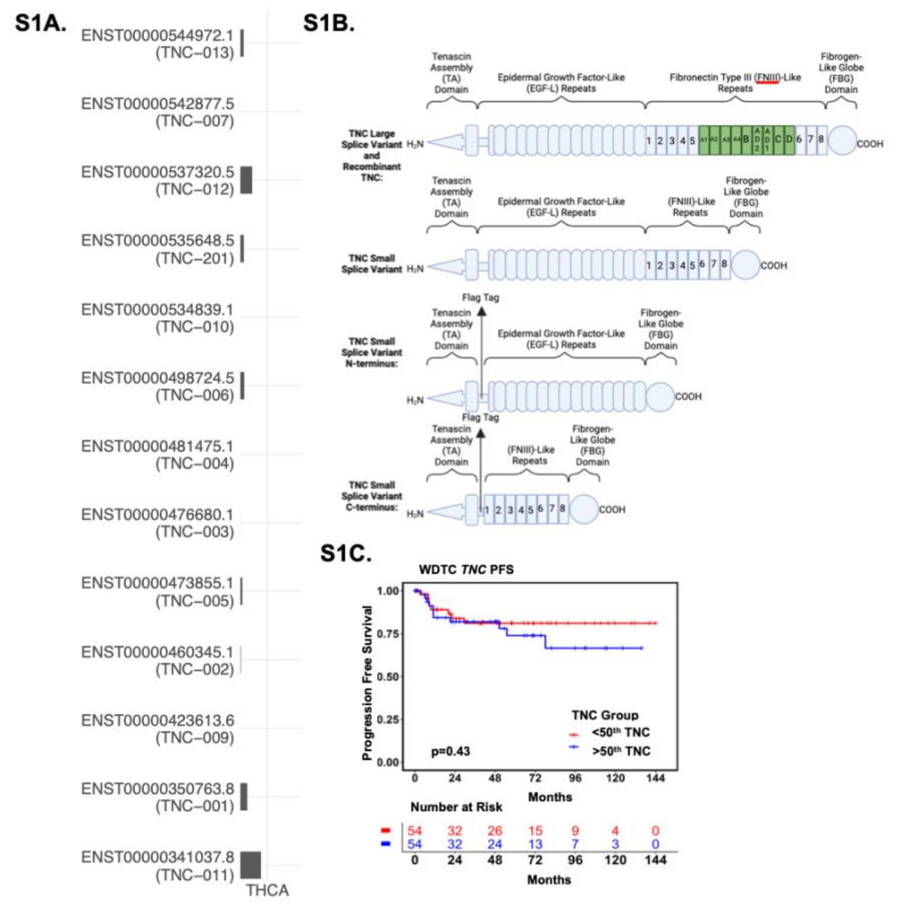


**Supplementary Figure 1.**
**(A)** GEPIA expression of several TNC variants present in thyroid cancer. **(B)** Schematic of TNC structure. Figure created using BioRender. **(C)** Progression-free survival plot for all malignant patients with less than 50th percentile *TNC* expression (red) or greater than 50th percentile TNC expression (blue) in primary tumors. p-values calculated using log-rank test.

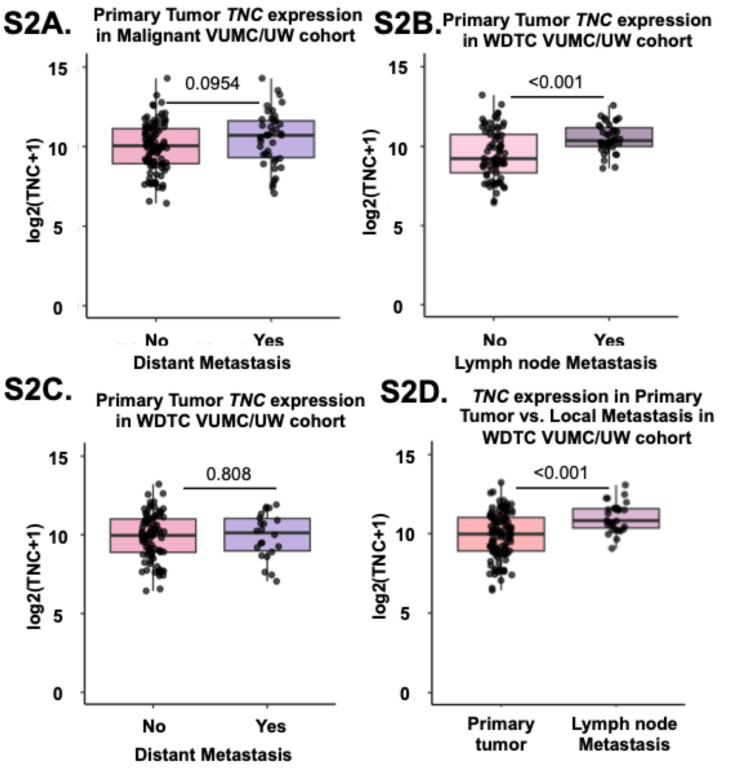


**Supplementary Figure 2.**
**(A)** Boxplots showing TNC expression in VUMC/UW primary malignant tumors split by presence (YES) or absence (NO) of associated distant metastases. p-values calculated using Wilcoxon rank-sum test with Bonferroni correction. **(B-C)** Boxplots showing TNC expression in VUMC/UW primary WDTCs split by the presence (YES) or absence (NO) of associated **(B)** local metastases or **(C)** distant metastases. p-values calculated using Wilcoxon rank-sum test with Bonferroni correction. **(D)** Boxplots showing TNC expression in VUMC/UW WDTCs split by primary tumor versus lymph node metastasis. p-values calculated using Wilcoxon rank-sum test with Bonferroni correction.

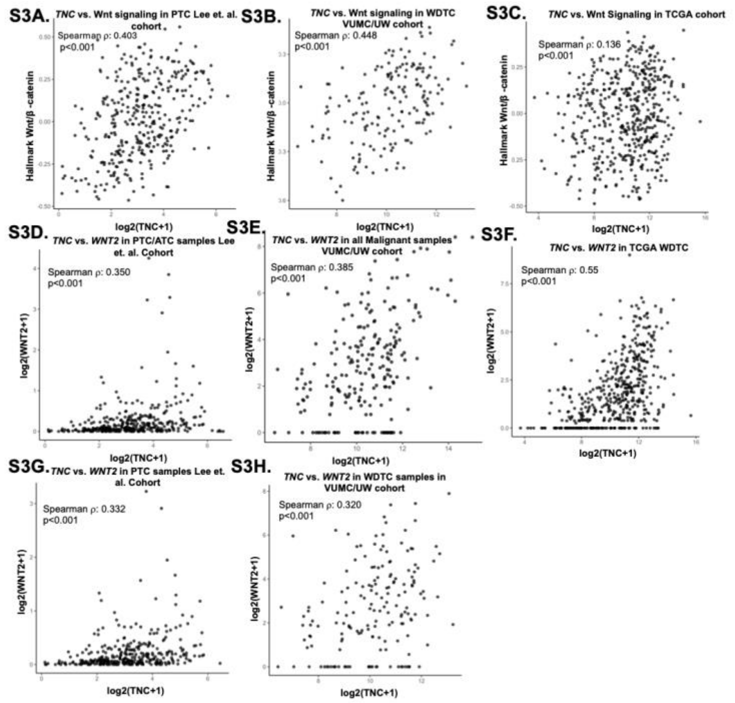


**Supplementary Figure 3.**
**(A-C)** Spearman's correlation between TNC expression and hallmark Wnt/β-catenin gene activity in **(A)** Lee et. al. WDTCs, **(B)** VUMC/UW WDTCs, and **(C)** TCGA WDTCs. **(D-H)** Spearman's correlation between TNC and Wnt-2 expression in **(D)** Lee et. al. malignant cohort, **(E)** VUMC/UW malignant cohort, **(F)** TCGA WDTCs, **(G)** Lee et al. WDTCs, and **(H)** VUMC/UW WDTCs.

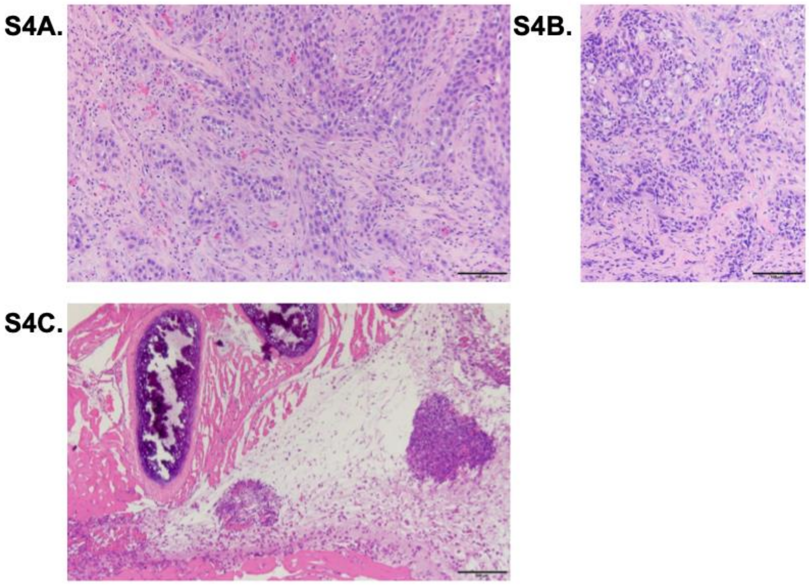


**Supplementary Figure 4.**
**(A)** H&E staining of an ATC patient tumor that had TNC-expressing leading-edge tumor cells. **(B)** H&E of 16T-PDX mouse tumor. **(C)** H&E staining of TNC 16T-PDX tumor invading into the chest wall.
